# Domestication of the ancient grain *Amaranthus cruentus* L

**DOI:** 10.1101/2025.11.27.690906

**Authors:** Roger Antonio Sulub-Tun, Akanksha Singh, Rolando Cifuentes-Velásquez, Markus G. Stetter, Ivonne Sánchez-del Pino

## Abstract

Grain amaranths are nutritious pseudocereals that have been domesticated in the Andes and North America mainly in Mexico. The repeated domestication of grain amaranths has recently been studied using genome-wide data. The results suggested independent processes of domestication from *Amaranthus hybridus* L. in different geographical regions. Although domestication process of the three grain amaranths has been partially elucidated, the detailed domestication centers remain unknown. Therefore, we study the geographic origin and center of domestication of *A. cruentus* within North America. We reveal a high genetic diversity and distinct history, using whole genome sequencing of 138 accessions belonging to *A. cruentus* and its closely related species (*A. hypochondriacus* L.) along with the wild relative *A. hybridus* mainly from South of Mexico and Guatemala. *A. cruentus* was most likely domesticated in central Mexico and then taken to Guatemala, where this species is most widely preferred and cultivated today. We identified three genomic groups of *A. cruentus*. *Amaranthus cruentus* group 1, representing the center of Mexico, *A. cruentus* group 2 mostly from Guatemala, and *A. cruentus* group 3 including individuals in Mexico, but distinct from group 1. Compared to *A. hybridus*, *A. cruentus* group 3 exhibited the lowest genetic differentiation and the highest genetic diversity among the *A. cruentus* groups. We find gene flow between *A. cruentus* group 1 and *A. hybridus* indicating the continuing interbreeding between wild and domesticated amaranth in its domestication center. Our results support the hypothesis of a Mexican origin of *A. cruentus* and a secondary center of domestication in Guatemala.

## INTRODUCTION

*Amaranthus* L. (Amaranthaceae) includes around 70 species, of which 40 are native from America (Berghofer & Schoenlechner, 2002). The genus includes species widely used as potherbs, ornamental plants, and for medicinal purposes (Bhandari, 1938). Multiple species of amaranth also grow as unwanted weeds leading to high yield losses (Steckel, 2007; Das, 2016). The genus also includes species known as the pseudocereal grain amaranths, which are a valuable source of seeds with high nutritional value (Steckel, 2007; Das, 2016). *Amaranthus* species are dicots and use C_4_ photosynthesis pathway (Sauer, 1993), which allows them to grow rapidly even under drought and other challenging environmental conditions such as salinity, heat, and ultraviolet irradiation (Omami & Hammes, 2006; Jamalluddin *et al*., 2019).

Grain amaranths comprise three species (*A. caudatus* L., *A. cruentus*, and *A. hypochondriacus*) that have been domesticated in different regions of the Americas. The grain amaranths are part of an ancient crop from “Central America” (*A. cruentus*, and *A. hypochondriacus*) and South America (*A. caudatus*). Early works identified a group of species that included the three grain amaranths and their putative wild relatives based on qualitative character such as, seed color, bract to utricle length ratios, and tepal posture (Sauer, 1950, 1967). The three domesticated species of *Amaranthus*, together with *A. hybridus* and *A. quitensis* Kunth form a monophyletic group and are part of the called “hybridus-species complex” (Kietlinski *et al*. 2013), which circumscription as a group of five species, has been the most followed in literature (e.g. Adhikary & Pratt, 2015; Stetter & Schmid, 2017) Floral variation in the hybridus-species complex consists of quantitative differences between minute flower parts, complicating the morphological distinction between species within the complex (Kietlinski *et al*., 2013).

Morphological works provided the basis to propose hypotheses to explain the origin of the three domesticated species, including the involvement of additional wild relatives (Sauer, 1950, 1967). Sauer (1967) proposed that distinct wild *A. hybridus* lineages served as progenitors for the domesticated taxa in North and South America. Genomic and phylogenetic analyses have demonstrated that *A. hybridus* is polyphyletic, comprising genetically distinct North and South American lineages that each contributed differently to the domesticated grain amaranths (Stetter & Schmid, 2017; Thapa *et al*., 2021; Gonçalves-Dias *et al*., 2023). Post-domestication gene flow was observed between the Mexican *A. hypochondriacus*, and the South American *A. caudatus* but only restricted gene flow between *A. hypochondriacus* and *A. cruentus*, despite their geographic co-occurrence (Gonçalves-Dias *et al*., 2023). Together, this evidence supports the scenario of multiple, regionally independent domestications of grain amaranths from polyphyletic *A. hybridus*.

*Amaranthus cruentus* is known as cahal scul, bledo, bledo Colorado, Xtes, among other vernacular names (Ibarra-Morales *et al*., 2021). Its seeds are eaten like cereals and the young leaves as a leafy vegetable (Das, 2016), representing an important element in the diet of local communities, such as the Otomies and the indigenous people from the Sierra Norte in Puebla, Mexico (Balcázar-Quiñones *et al*., 2020; Mapes *et al*., 1996). Red inflorescence water extracts were used to make bread mixed with maize for ceremonial dances (Sauer, 1967). The species is also extensively used in Africa as leafy vegetable (Das, 2016).

*Amaranthus cruentus* most likely originated in “Central America”, where it is historically distributed and is recently cultivated, showing its wide distribution in various environments and climates (Grubben, 1967; Sauer, 1967; Grubben & Sloten, 1981). This crop is short-day species, flowering and setting seed when the day length is less than 12 h (Joshi et al., 2018), and is the most photoperiod-insensitive among grain amaranths (Brenner et al., 2000). This grain species is morphologically diverse, and can hybridize readily with wild relatives of *Amaranthus* (Makinde *et al*., 2010). It grows up to 2 m tall, has abundant flowers that form spikes (dichasial, cymose branching inflorescence units) in a long and dense terminal thyrsus (compound inflorescences consisting of thyrsoid paracladia) and is prickly due to acute tepals with pungent apices (Wolosik & Markowska, 2019). Its inflorescences present a variety of colors such as green, yellow, red, purple, etc., and they can reach up to 50 cm long and produce more than 50,000 seeds per inflorescence (Robertson & Clemants, 2003; Grubben, 2004). The ‘Blood Amaranth’ and ‘Kerala Red’ are cultivars of *A. cruentus* used as ornamental mainly due to its red color (Brenner & Makus, 1997; Das, 2016). Despite its high variation, wide distribution and local importance, little is known about the population structure and history of *A. cruentus* within its domestication center.

Domestication is an evolutionary process that favored plant traits of social, economic, cultural, ecological, and technological importance to humans leading to a set of traits that distinguish domesticated from the wild plants, known as domestication syndrome (Hammer, 1984). In many crops this process led to a loss of genetic diversity, retaining only a fraction of the original genetic diversity found in their wild relatives (Pickersgill, 2007; Chávez-Pesqueira, 2017). The region between Mexico and Guatemala has been described as one of the eight centers of plant domestication, and is defined by a high diversity in crops and their wild relatives (Vavilov, 1992). Key crops were domesticated here, including maize around 9000 calendrical years ago (cal. B.P.) (Piperno *et al*., 2009), common bean around 8000 years before present (YBP) (Mamidi *et al*., 2011) and squashes between 8000–10,000 cal. B.P. (Smith, 1997). Hence, anthropological research in the region between Mexico and Guatemala is required to understand the history of agriculture in early human civilizations.

For the grain amaranth *A. cruentus,* Sauer (1967, pg. 123), reported that “Amaranthus cruentus *evidently originated as a domesticated grain crop somewhere in southern Mexico or Guatemala, the only region where it is found in aboriginal cultivation within the range of its probable progenitor*, A. hybridus”. Yet, the exact domestication sites remain undetermined. In Guatemala and South of Mexico, *A. cruentus* has many uses, suggesting that domestication might have occurred there before spreading northward to Mexico (Williams & Brenner, 1995, cited in Espitia-Rangel *et al*., 2010). However, in Mexico, specifically in the Coxcatlán cave situated in the Tehuacán Valley, Puebla, remnants of several pre-Hispanic crops were discovered, including two species of amaranth, *A.* cf. *cruentus* L. and *A.* cf. *leucocarpus* (MacNeish, 1992), the latter now a synonym of *A. hypochondriacus* L. (Sauer, 1950). Seeds of *A. cruentus* found in the Coaxatlán cave were dated 4000 B.C, although their actual age may be even older (Sauer, 1993). Espitia-Rangel *et al*. (2010) concluded that the origin of *A. cruentus* was in “Mesoamerica”, close to the Mexican pre-Hispanic cultures of the biogeographic region of the Trans-Mexican volcanic Belt. Therefore, the exact domestication center and diversity of *A. cruentus* remains unknown. Genomic information of landraces and local wild relatives provide the opportunity to understand the origin and history of the crop.

In this study, we aim: 1) To test Sauer’s hypothesis about the center of domestication of *A. cruentus* postulated somewhere in south of Mexico or Guatemala by using whole genome sequencing (WGS) data, and 2) to propose the process of domestication of *A. cruentus* by analyzing intraspecific and interspecific (*A. hybridus* and *A. hypochondriacus*) patterns of genetic variation.

## MATERIALS AND METHODS

### Sampling and DNA extraction

The present study included 209 accessions of *A. cruentus* and its close relatives (*A. caudatus* = 37, *A. cruentus* = 81, *A. hybridus* = 26, *A. hypochondriacus* = 32, *A. quitensis* = 19, *A.*× *wallichii* = 1, Ambiguous = 8, hybrids = 5). Of these, 118 accessions were obtained as raw reads from Stetter *et al*. (2020) available in the Europe Nucleotide Achieve (ENA, https://www.ebi.ac.uk/ena/browser) under project number PRJEB30531. The remaining 91 accessions were grown from seeds obtained from multiple sources, including the United States Department of Agriculture (USDA), personal collections, and donations. Our sampling efforts were primarily focused on Mexico and Guatemala, although we also included samples from other countries of America (Supplementary Figure 1 and Supplementary Table 1). Material identification was followed based on names labeled for herbarium material, original sources such as GenBank names, key identifications in published papers, and data bases.

The seeds were germinated at the Yucatan Center of Scientific Research nursery. Subsequently, the plants were cultivated outdoor watered thrice weekly to facilitate optimal growth. Leaves from a single plant within each accession were carefully harvested and preserved in an ultra-freezer set to -80°C for long-term storage.

Approximately 100 mg of the plant material was crushed using mortar and pestle immersed in liquid nitrogen. DNA was extracted by using the CTAB extraction method (Doyle & Doyle, 1987), followed by subsequent incubation at 65 °C with 1 µL of RNase (Qiagen) for one hour. The integrity of the extracted DNA was verified by electrophoresis in a 0.8% agarose gel. The concentration and purity were determined by reading absorbances in a nanodrop.

### Libraries preparation and sequencing

The whole genome sequencing libraries were prepared using modified Nextera-shallow sequencing protocol as described in Rowan *et al*. (2019). First, we mixed 2 µL of DNA (conc. 0.5 ng/µL), 2.1 uL of autoclaved water, 0.818 µL tagmentation buffer and 0.0818 µL tagmentation enzyme from Illumina Tagment DNA Enzyme and Buffer Kit (Cat. No. 20034197). The tagmentation mix was incubated at 55^°^C for 10 minutes. Later on, it was cooled at room temperature until get ready to add 5uL of custom index-barcoding primers. The sample were amplified using KAPPA2G Robust PCR kit with GC buffer (Cat. No. KK5004). The PCR conditions were: 72^°^C for 3 min, 95°C for 1 min, followed by 14 cycles at 95°C for 10 s, 65°C for 20 s, and final extension at 72°C for 3 min. The library success and its size distribution were analyzed on 2% agarose gel. Furthermore, 5uL of each of the successful libraries were pooled and proceeded for dual size selection (range 300-500bp) using Promega Pronex-Size Selective Purification System following manufacturer’s instructions. After quality control, libraries were sequenced at 2 × 150 base pairs (bp) on an Illumina NovaSeq 6000 sequencer (Novogen, Germany).

### Single Nucleotide Polymorphism (SNP) calling and filtering

Quality assessment was performed on the raw reads using FastQC (Andrews, 2023). The results were summarized using MultiQC 1.14 (Ewels *et al*., 2016). Good quality (based on the Phred score) reads were mapped to the reference genome of *A. cruentus* (Ma *et al*., 2021) using BWA-MEM2 (Vasimuddin *et al*., 2019). The raw reads from Stetter *et al*. (2020) and two accessions of *A. tuberculatus* (ERR3220318 and SRR16300825) (Kreiner *et al*., 2019, 2023) were also mapped on the same reference genome.

Duplicates were marked using Picard (http://broadinstitute.github.io/picard), and flagstat files (the alignment statistics) were generated using SAMtools (Danecek *et al*., 2021). Only samples with high mapping percentage (>90 %) were included.

Single Polymorphism Nucleotide (SNP) were called using ANGSD: Analysis of next generation Sequencing Data (Korneliussen *et al*., 2014) with the following parameters: P 5, doCounts 1, doGeno 3, dopost 2, domajorminor 1, domaf 1, doBcf 1, snp_pval 1e-6, remove_bads 1, minMapQ 30, minQ 30, df 2, checkBamHeaders 0, minInd 47, only_proper_pairs 1, trim 0, setMinDepth 47, setMaxDepthInd 150. The variants were filtered using BCFtools (Danecek *et al*., 2021) for a minimum allele frequency (MAF) of 0.01 and only variants with less than 10% missing genotypes were included.

### Genetic determination

Since morphology-based identification is complicated in amaranth (Costea *et al*., 2006; Singh & Stetter, 2025), molecular analysis was further used to confirm the taxonomic identifications by first looking at population subdivision and the relationship of *A. cruentus* within the hybridus-species complex. This helped to confirm or re-assign the taxonomic identification of each sample based on molecular instead of morphological criterium. A Principal Component Analysis (PCA) was carried out using PCAngsd (Meisner & Albrechtsen, 2018) based on genotype likelihoods from ANGSD. In order to confirm the morphological identification of each sample, we included the three domesticated species as well as the wild *A. hybridus* and *A. quitensis*, considering species identification based on Stetter *et al*. (2020) and Gonçalves-Dias & Stetter (2021).

### Genetic structure and genetic diversity

To study the genetic and population structure, we further used *A. cruentus* (96), *A. hybridus* (12) and *A. hypochondriacus* (30) accessions (see Supplementary Table 1 for more information). We analyzed population structure performing new PCA with the samples previously listed. We also used NGSadmix (Skotte *et al*., 2013) for values from K = 2 to K = 12 to identify the most optimal number that describes the genetic structure of the study sampling. Weir and Cockerham F_ST_ estimates were calculated with Vcftools (Danecek *et al*., 2011) among the populations identified with NGSadmix. Nucleotide diversity (π) was also estimated for these populations using ANGSD. We created a Neighbor Joining tree, including two accessions of *A. tuberculatus* as outgroup using the ape (Paradis & Schliep, 2019) and adegenet (Jombart & Ahmed, 2011) R libraries. For this tree we LD-pruned the filtered SNPs using PLINK2 (Chang *et al*., 2015) with a window size of 50 SNPs and a R^2^ value of 0.3 and keeping only SNPs with less than 2% missing values. This resulted in a set of 433,630 SNPs to construct the tree.

### Estimating Effective Migration Surfaces

The genetic similarity decays with geographic distance (isolation by distance), the visualization of this pattern is possible using effective migration surfaces, which illustrate spatial variation in effective migration. In this context, effective migration is interpreted as an index of relative gene flow: lower values indicate regions where genetic similarity decays rapidly with distance (potential barriers to gene flow), while higher values indicate regions of increased connectivity (Petkova *et al*., 2016). Therefore, we conducted FEEMS (Fast Estimation of Effective Migration Surfaces) (Marcus *et al*., 2021) to estimate migration surfaces for *A. cruentus* across Mexico and Guatemala.

FEEMS requires three data sets to be performed: 1) the sampling coordinates, 2) a triangle grid (in this study it was created using ArcGIS with 67.114 km), and 3) the genetic data in PLINK format. From the data set used for Neighbor Joining analysis, we extracted individuals of *A. cruentus* with geographical data information. We also extracted 10% of random SNPs. The resulting data set included 57 individuals (Supplementary Table 1) and 573,111 SNPs. Different values of lambda were evaluated and the optimal value was the one with the lowest cv l2 error.

### Introgression analysis

Gene flow was inferred estimating D-statistic by using ABBA-BABA2 function and implemented in ANGSD. D values were estimated between groups identified in NGSadmix analysis with k = 5. Two accessions of *A. tuberculatus* (ERR3220318 and SRR16300825) (Kreiner *et al*., 2019) were used as outgroup. To test geographical origin of genetic groups and based on the fact that *A. hybridus* is polyphyletic, we performed ABBA-BABA2 by grouping *A. hybridus* in three different arrangements: 1) *A. hybridus* as a single group (12 individuals) wide distributed in America, 2) *A. hybridus* from Mexico + Central America (countries connecting North and South America: Belize, Guatemala, El Salvador, Honduras, Nicaragua, Costa Rica, and Panama) (6 individuals), different of *A. hybridus* from South America (5 individuals), and 3) *A. hybridus* from Mexico (1 individual) different of *A. hybridus* from Guatemala (5 individuals).

## RESULTS

### Reads and mapping quality

The 91 newly sequenced samples coverage ranged from 2.2 and 15.6 x, with an average mapping rate of 96.94% ± 1.31% on the reference genome of *A. cruentus* (Ma *et al*., 2021). We also mapped 118 accessions of *A. cruentus* and relatives taken of Stetter *et al*. (2020) with the reference genome of the same species, which resulted in 94.71% ± 1.26% of reads correctly matched. Five individuals were discarded due to low mapping percentage (Supplementary Table 1). We obtained a total of 42,862,466 variants, and 23,862,427 biallelic SNPs after filtering.

### Taxonomic determination based on molecular data

The principal component analysis (PCA) showed strong population structure, with the first two components explaining together 95.43% of variation in the data. PC1 and PC2 separated the study samples in four groups, one group included all *A. cruentus* accessions, a second group included all *A. hypochondriacus* samples, a third group included together the accessions of *A. caudatus* and *A. quitensis*, and in the center, a fourth group was clustering the individuals of *A. hybridus* (Figure 1). We identified 12 individuals from previous studies, as well as 21 new accessions included in this study where species morphologically assignment did not fit with the genetic clustering using PCA (Figure 1 and Supplementary Table 2).

**Figure 1.**
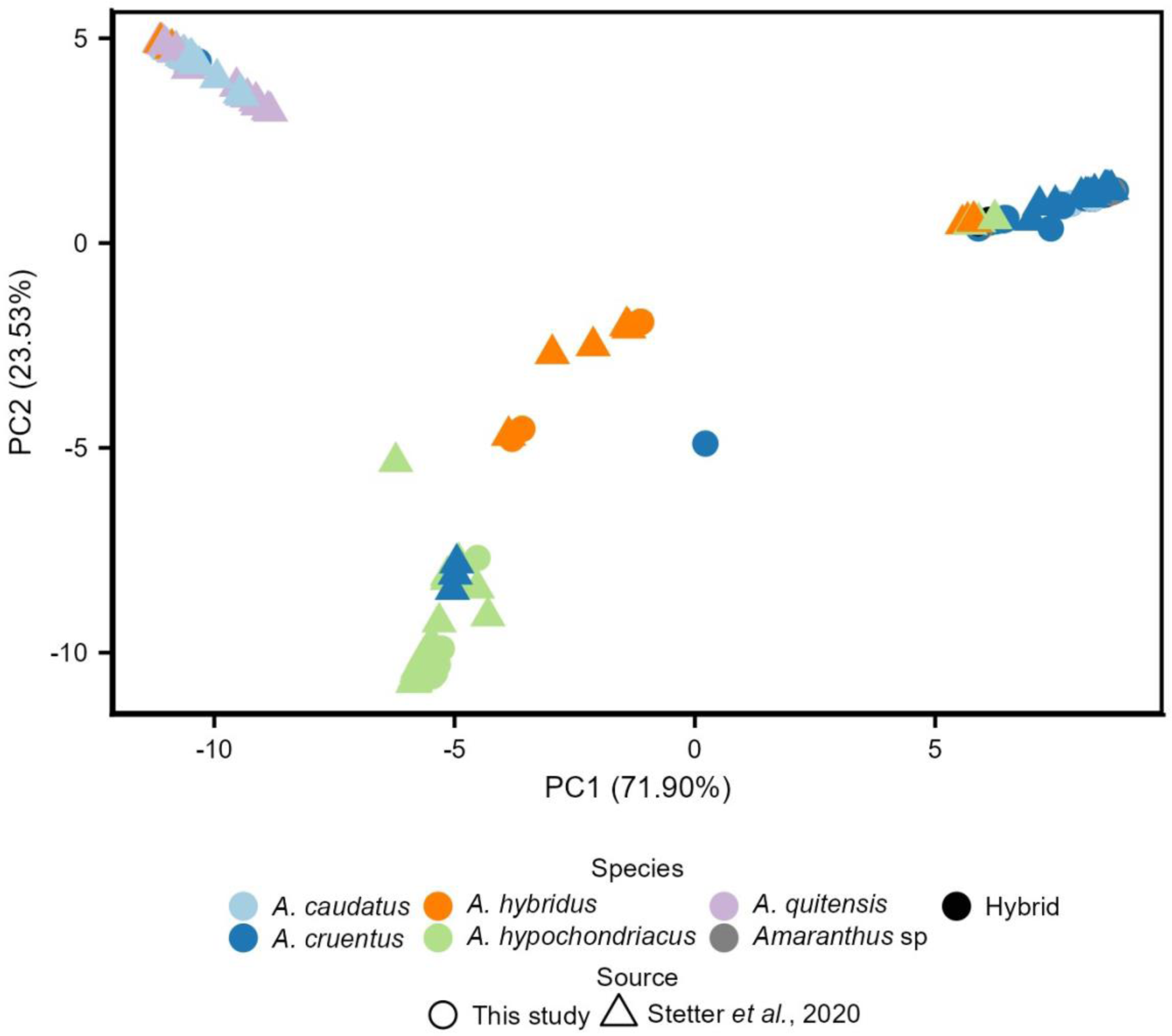
Plot of the first two axes from principal component analysis including the three grain amaranths, two wild relatives (*A. hybridus* and *A. quitensis*), and other *Amaranthus* taxa.

### Genetic diversity and structure reveal three genomic groups of *A. cruentus*

The population genetic analyses revealed intraspecific patterns of genetic variation within *A. cruentus*. In the PCA including only *A. cruentus*, *A. hybridus*, and *A. hypochondriacus* (after species reassignment), the first two principal components explained together 93.41% of variation and separated the studies samples in three groups; one group included all *A. cruentus* accessions, a second group included all *A. hypochondriacus* samples, and a third group included the accessions belonging to *A. hybridus* (Figure 2A).

**Figure 2.**
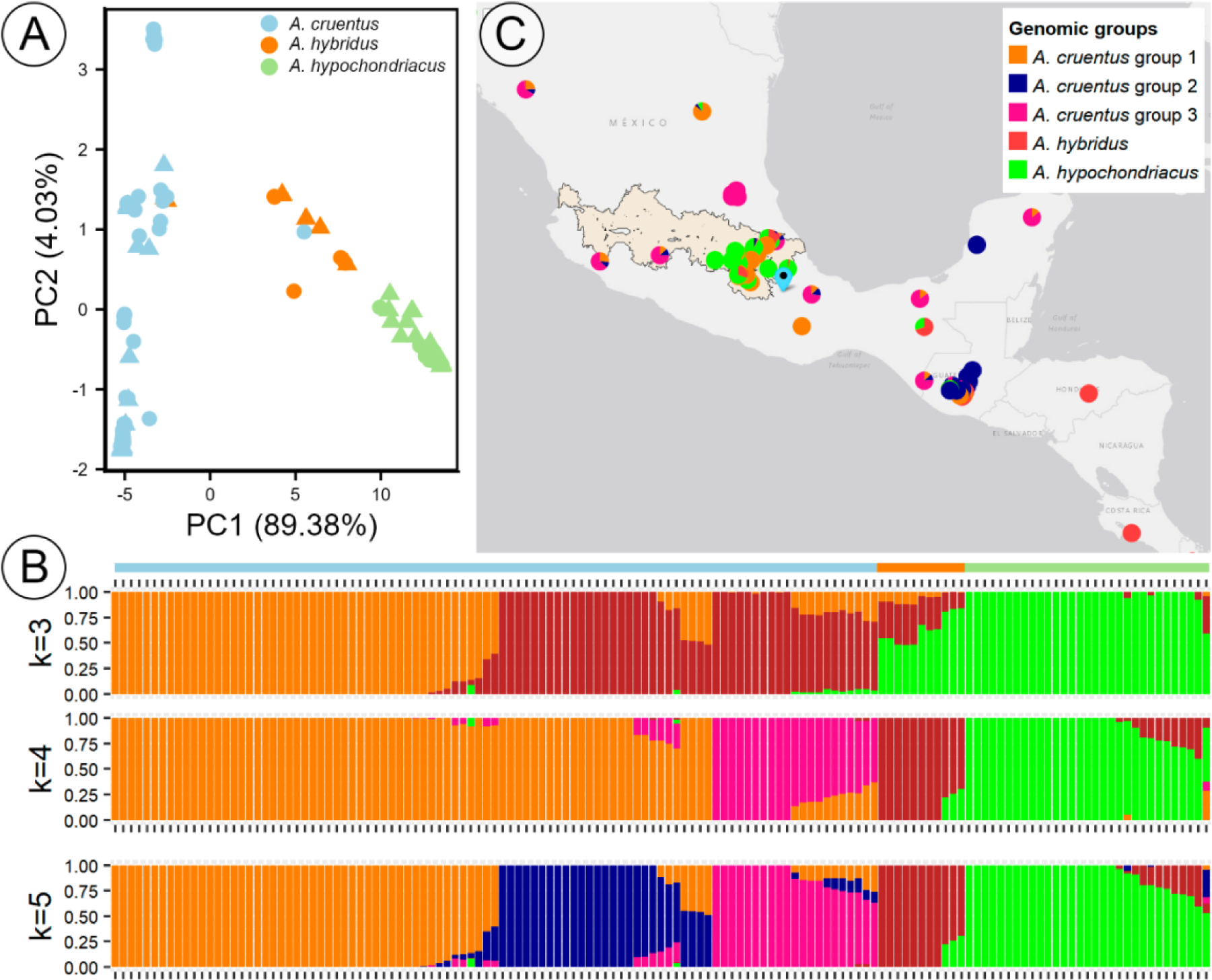
Genetic structure analyses and geographical distribution of *A. cruentus* and relatives. A) Plot of the first two axes from principal component analysis, circles represent new sequenced samples and triangles samples of Stetter *et al*.’s (2020) study. B) Admixture proportions plots showing from k = 3 to k = 5, bar colors above represent species of *Amaranthus* based on the principal component analysis. C) geographical distribution of sampling from Mexico and Central America, pie charts represent admixture proportions based on k = 5. Light orange area represents the Trans-Mexican volcanic Belt, and the cyan flag indicate the location of the Coxcatlán cave.

The cross validation based on Elbow method for NGSAdmix indicated that the optimum K is 5 (Supplementary figure 2). When three groups are assumed, we observed a separation of *A. cruentus* into two groups, with a mix of *A. hybridus* and *A. hypochondriacus*, suggesting a closer relationship compared to *A. cruentus*. At K = 4, there is a clear separation of *A. hybridus* from the two domesticated species (*A. cruentus* and *A. hypochondriacus*). With a K of five, three groups were resolved within *A. cruentus* of which, one is mostly a Guatemalan group and two are Mexican groups (Figures 2B and 2C).

The Neighbor Joining tree confirms the separation of *A. cruentus* into three main clades, similar to the admixture analysis. *Amaranthus hypochondriacus* individuals were grouped in a single clade. *Amaranthus hybridus* was polyphyletic with some representatives embedded within the *A. cruentus* and *A. hypochondriacus* clades.

### Patterns of genetic differentiation and diversity provide further insights into the domestication history of *A. cruentus*

When we compared the F_ST_ estimations between the *A. cruentus* groups and *A. hybridus*, we found a lower genetic differentiation between *A. cruentus* group 3 and *A. hybridus* (F_ST_ = 0.35); followed by *A. hybridus* and *A. cruentus* group 2 (F_ST_ = 0.49), and the higher value was found between *A. hybridus* and *A. cruentus* group 1 (F_ST_ = 0.56) (Figure 3A). The genetic differentiation between *A. hypochondriacus* and *A. hybridus* is lower compared with any of the genomic groups characterized in *A. cruentus* (F_ST_ = 0.31) (Figure 4A).

**Figure 3.**
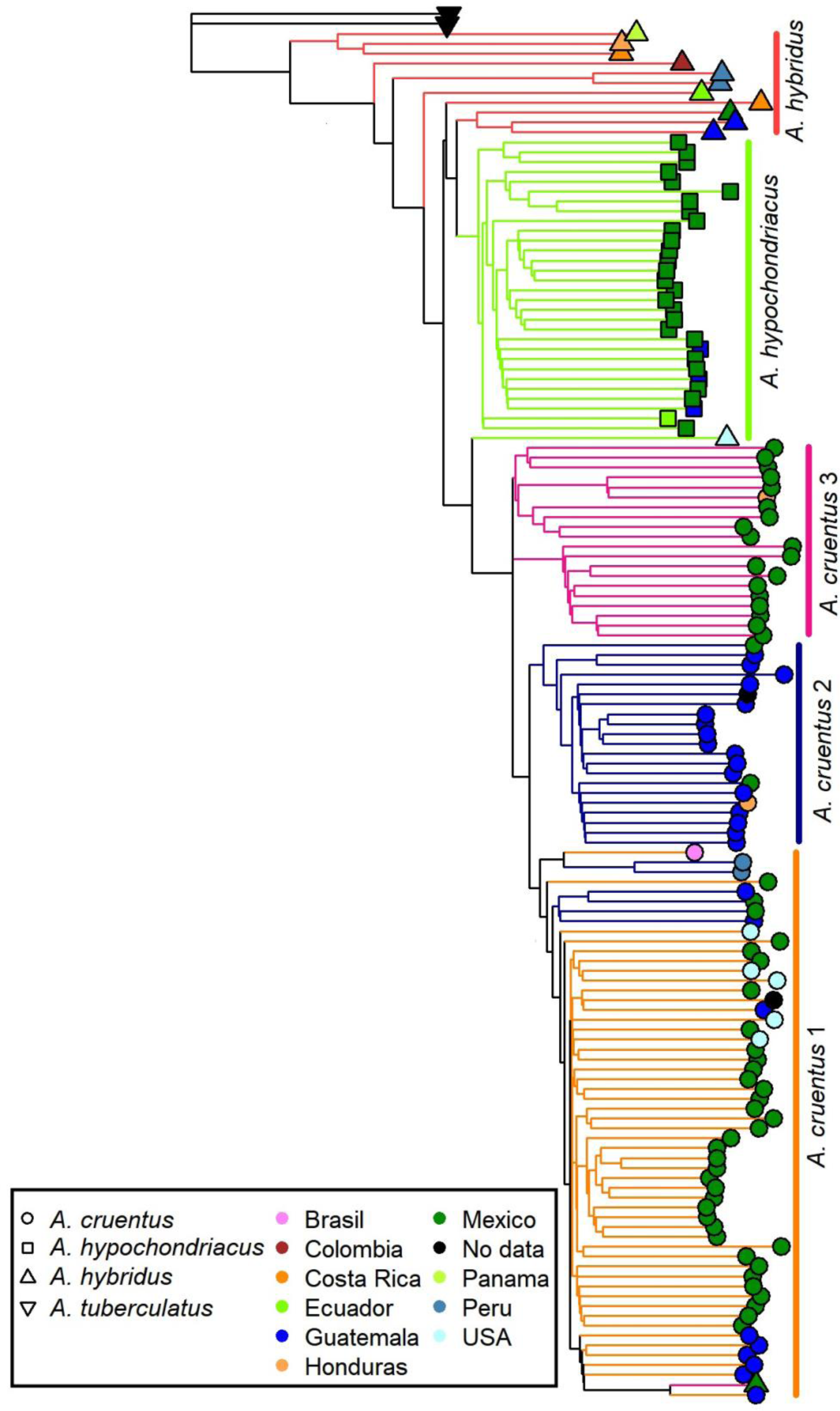
Neighbor Joining tree based on 433,630 SNPs. Colors represent countries of origin, ND = no data. Color of the branches represent the five genomic groups identified in Admixture analysis with K = 5.

**Figure 4.**
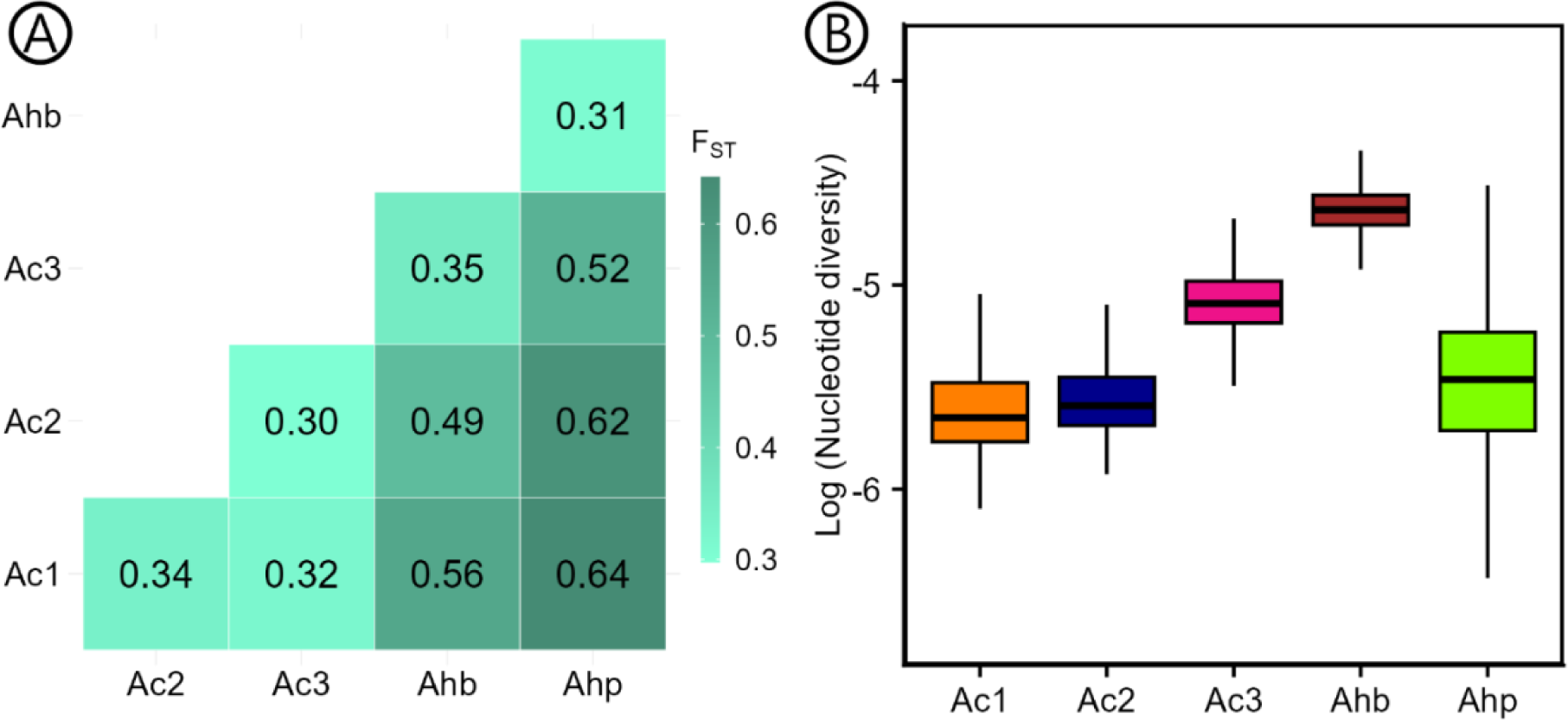
Genetic differentiation and nucleotide diversity of *Amaranthus*. A) Pairwise F_ST_ between populations identified with k = 5. B) Logarithm of the nucleotide diversity (π) of the genomic groups of *A. cruentus* based on k = 5 estimated over 50 kb windows on the whole genome. Ac1: *A. cruentus* group 1, Ac2: *A. cruentus* group 2, Ac3: *A. cruentus* group 3, Ahb: *A. hybridus*, Ahp: *A. hypochondriacus.*

We further estimated genetic diversity based on the above defined groups. *A. hybridus* showed the highest average nucleotide diversity (log(π) = -4.634). Among the three genomic groups of *A. cruentus*, the group 3 had the highest nucleotide diversity (log(π) = -5.0786). *Amaranthus cruentus* groups 1 (log (π) = -5.5971) and 2 (log (π) = -5.5470), and *A. hypochondriacus* (log (π) = -5.4779) exhibited similar nucleotide diversity (Figure 4B).

### *Amaranthus cruentus* migrations surfaces identifies geographic patterns of diversity

To study the geographical pattern of diversity in *A. cruentus,* we estimated the migration surface using FEEMS (Marcus *et al*., 2021). Based on cross-validation, the optimal smoothing value for λ is 0.00616 (cv l2 error = 0.08527). The estimated migration surface revealed higher effective migration rates in central Mexico (log10(w) > 1), whereas areas of reduced effective migration (log10(w) < 1) were evident in southern Mexico and Central America, reflecting topographic or ecological barriers such as the Isthmus of Tehuantepec or mountainous regions such as the Sierra Madre Occidental, Sierra Madre Oriental, and Sierra Madre del Sur (Figure 5).

**Figure 5.**
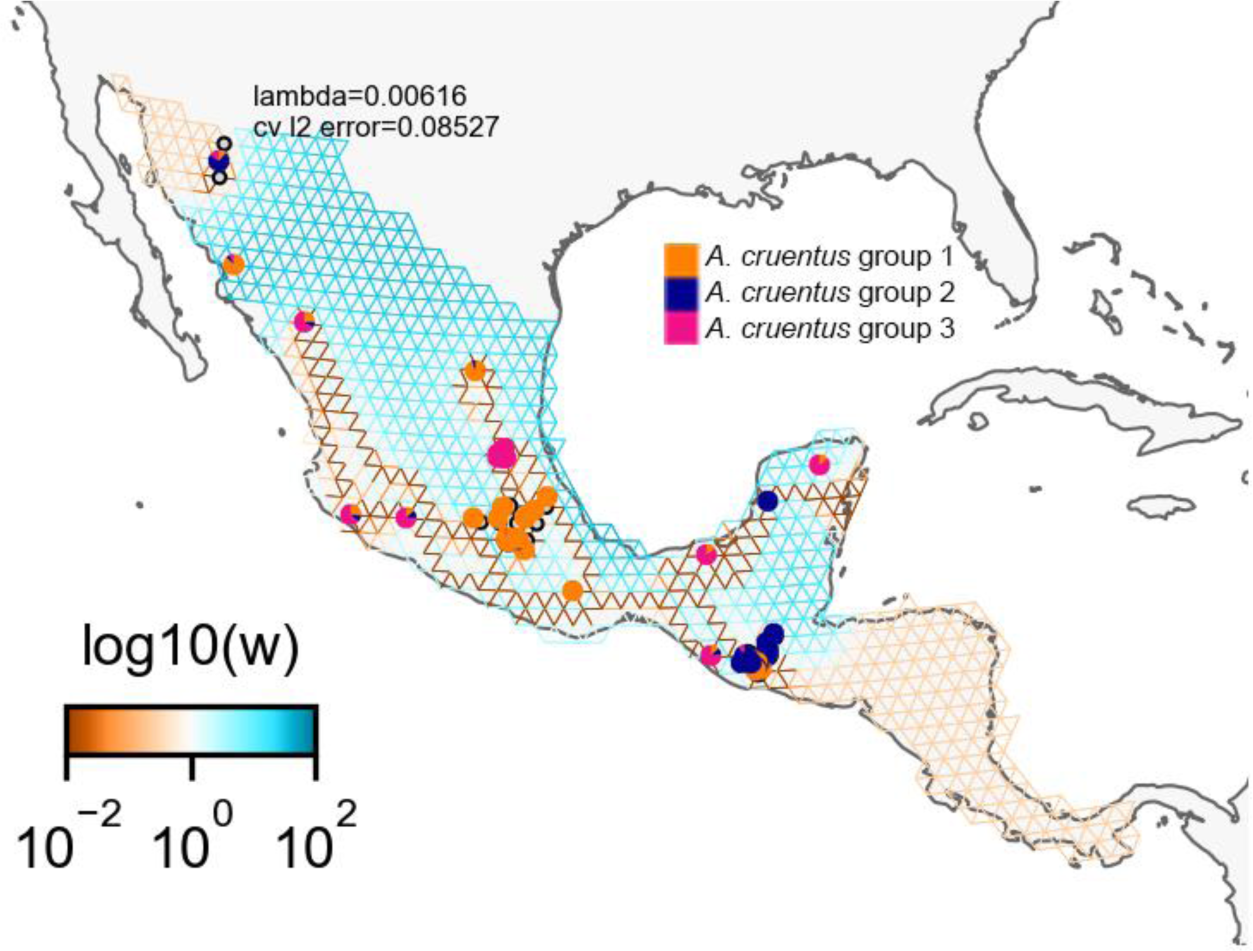
FEEMS map of *Amaranthus cruentus* from Mexico and Guatemala. Blue highlighted regions represent higher than expected migration rates and red highlighted regions lower than expected migration rates.

### Introgression between wild and domesticated amaranth shaped crop diversity

We investigate the level of gene flow using an ABBA-BABA test. This analysis proposed different results based on three applied assumptions. First, when *A. hybridus* is considered as a single group, then no significant values for D arose between *A. hybridus* and *A. cruentus* (Figure 6A), meaning that alleles shared between the species are ancestral. Second, when *A. hybridus* was divided into 1) Mexican + Central American and 2) South American subgroups, significant introgression values were detected between all three groups of *A. cruentus* and the two subgroups of *A. hybridus*. However, D values were higher between groups of *A. cruentus* and *A. hybridus* from Central America (Figure 6B). Third, when Mexican + Central American *A. hybridus* were divided into Mexican and Guatemalan subgroups, introgression was detected between *A. cruentus* group 1 and *A. hybridus* from Mexico (D = 0.2973), and between *A. cruentus* group 3 and *A. hybridus* from Guatemala (D = 0.0197); no significant D values were found between *A. cruentus* group 2 and any groups of *A. hybridus* (Figure 6C). In all three analyses, higher D values were detected among *A. cruentus* groups, while lower D values were estimated among *A. cruentus* and other species (i.e., *A. hybridus* and *A. hypochondriacus*), showing stronger gene flow within a crop species than between crop species.

**Figure 6.**
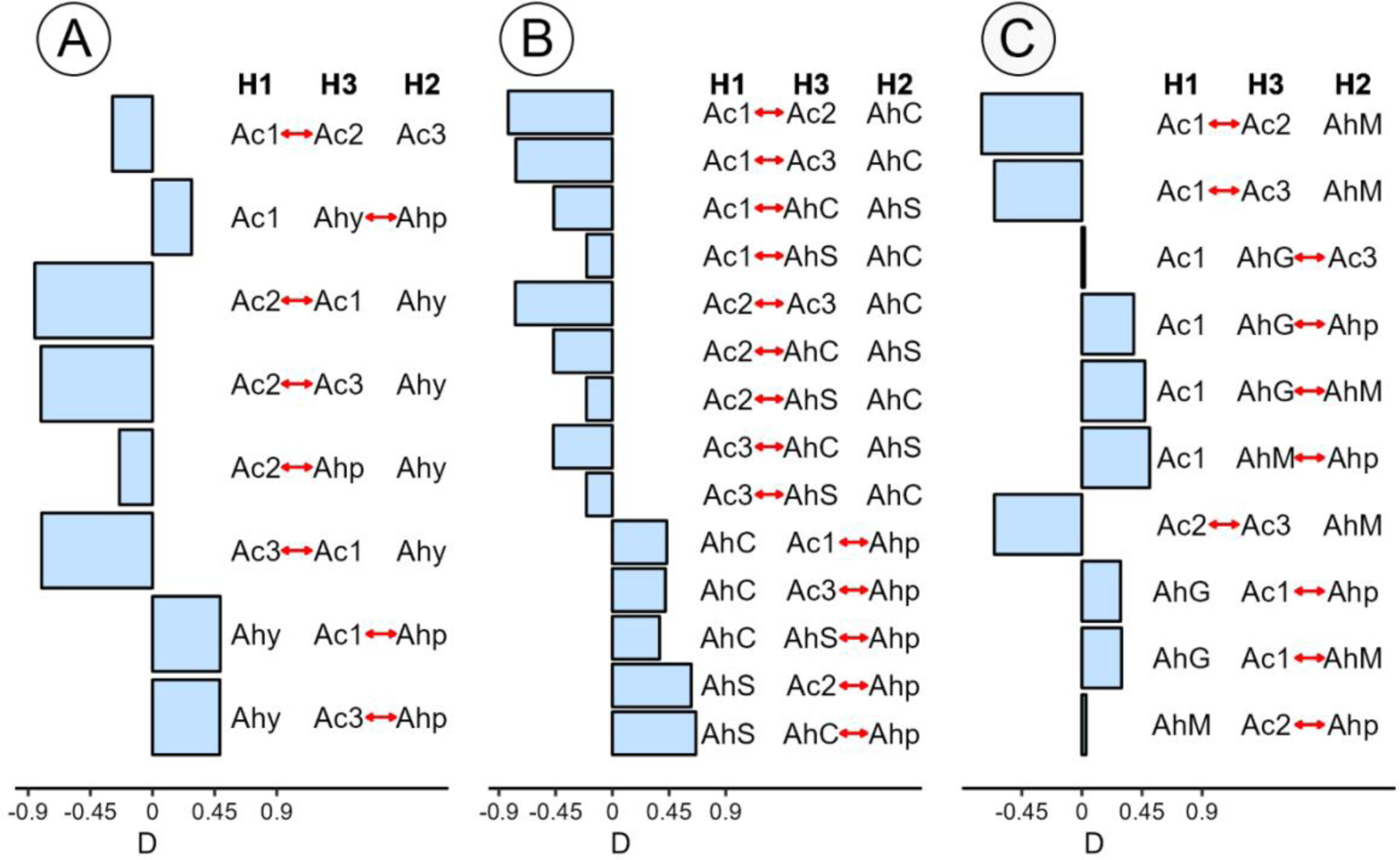
D statistics among populations of species of *Amaranthus* evaluated in this study. A) *A. hybridus* was considered as single group; B) *A. hybridus* was separated into 1) Mexican + Central American and 2) South American accessions; C) Mexican + Central American populations of *A. hybridus* separated into Mexican and Guatemalan accessions. Ac1 = *A. cruentus* group 1, Ac2 = *A. cruentus* group 2, Ac3 = *A. cruentus* group 3, Ahp = *A. hypochondriacus*, Ahy = *A. hybridus*, AhS = *A. hybridus* South America, AhC = *A. hybridus* Mexico + Central America, AhM = *A. hybridus* Mexico, AhG = *A. hybridus* Guatemala. Only significant non redundant trees (absolute value of z score ≥ 3) were included. Red arrows denote gene flow.

## DISCUSSION

### Historical records of *Amaranthus cruentus*

The importance of *A. cruentus* for the subsistence of pre-Columbian civilizations is undeniable, not only as part of the diet of ancient people but also for religious purposes (Smith, 1961). Archaeological evidence shows that it has been cultivated since early agricultural civilizations in Tehuacán Valley as a staple crop with maize, beans, and squashes, supporting its role as an important crop for the diet and cosmovision of ancient societies (MacNeish, 1992; Sauer, 1993).

Chronology of the Tehuacán Valley has been traced based on the excavations made (for further information of other phases see MacNeish *et al*., 1967). At the beginning of the Coxcatlán phase, dated from 4500 to 3500 B.C., people were still plant collectors, using wild maize, chili, avocados and gourds, by the middle of this period they acquired amaranth, beans, squashes and others. At the end of this phase, the people probably were organized as macrobands (large communities) for longer periods in the valley, but they were not completely sedentary, and they broke up into microbands (small communities) again during dry seasons (MacNeish *et al*., 1967). However, Sauer (1969) indicated that specimens found in the Coxcaltán cave do not show transitional stages from wild to cultivated, and so, there is no evidence of local domestication of grain amaranths in the Tehuacan Valley. Thus, the seeds of the Coxcatlán cave could be domesticated in other place and transported to there (Sánchez-del Pino *et al*., 2025)

*Amaranthus cruentus* held significant cultural and ritual ceremonies in pre-Columbian civilizations in addition to its nutritional importance. The Aztecs for example, used the seeds to create figures called *tzoalli*, which were consumed during ceremonial rituals (Sauer, 1993). For this reason, during the Spanish conquest the cultivation of *A. cruentus* was forbidden, but it persisted in some rural areas and recently resurged due to its high nutritional value and resilience (Brenner *et al*., 2000).

### The center of domestication of *Amaranthus cruentus*

Despite their population structure and high genetic differentiation, the amaranth populations can well interbreed. Historical interbreeding between populations would have led to gene flow among the groups. Previous work has shown strong but distinct patterns of introgression between grain amaranth species in South and North America. Our work confirms the restricted gene-flow between *A. cruentus* and the sympatric *A. hypochondriacus* (Gonçalves-Dias *et al*., 2023). Our D-statistic additionally identified significant introgression between *A. cruentus* group 1 and *A. hybridus* and between *A. cruentus* group 3 and *A. hybridus*. In addition, we did no detect significant introgression between the group from Guatemala and *A. hybridus* (Figure 6C). *Amaranthus cruentus* group 1 is mainly located in the Trans-Mexican volcanic Belt, close to the Coxcatlán cave; *A. cruentus* group 3 is mainly located northward, in the state of San Luis Potosí and some scattered individuals distributed along Mexico (Figure 2 C), this group is also the most genetically similar to *A. hybridus* (Figure 4A). According to these findings, *A. cruentus* might have been domesticated in Mexico, based on the assumption that the genetic pool is similar to the wild ancestor and hence supporting the theory of the Mexican origin proposed by Espitia-Rangel *et al*. (2010) based on geographical distribution of *Amaranthus* accessions collected in Mexico. The finding also aligns with the oldest archaeological remains in Mexico found in the Coxcatlán cave of *A. cruentus* (MacNeish, 1992). Even if the found seeds were not domesticated there, here we suggest that this process probably may have taken place elsewhere, likely within the central region of Mexico.

Hence, our results suggest that Mexico could be considered the primary center of domestication. Genetic and phenotypic variations in *A. cruentus* found in Guatemala, in addition to environmental and cultural differences between central Mexico and Guatemala may have driven further diversification after the crop spread southward and arose from post-domestication dispersal and adaptation to local conditions (Sauer, 1993; Espitia-Rangel *et al*., 2010).

The resulting migration surface revealed regions of elevated effective migration concentrated in central Mexico, suggesting historical or ongoing gene flow through these areas, potentially reflecting environmental continuity or seed exchange routes of *A. cruentus* (Figure 5). On the other hand, gene flow barriers were detected in southern Mexico and Central America, potentially corresponding to ecological or topographic barriers such as the Isthmus of Tehuantepec and surrounding mountainous regions, reducing gene flow and promoting differentiation (Figure 5). These findings underscore how geographic barriers detected in this study, human cultural practices (Kietlinski *et al*., 2013), and multiple domestications (Sauer, 1967; Mallory *et al*., 2008; Stetter., 2020) have jointly shaped the genomic diversity of *A. cruentus*.

We cannot exactly say when or how amaranth arrived in Guatemala, but there is evidence of latter influence of Mexican cultures on Guatemalan people. For example, it is suggested that people from Teotihuacán play a role in the formation of Kaminaljuyú, one of the historically most important sites in Mesoamerica and located in the Valley of Guatemala (Sande rs *et al*., 1974). Teotihuacán and Kaminaljuyú are places separated by more than 1,000 km; however, they were connected via a complex web of social and economic relations (Ferguson, 2007; Arroyo *et al*., 2020). Another example is the Olmec legacy on of Cival in northeastern Petén, Guatemala (Estrada-Belli, 2006). We suggest that his connection might have led to gene flow between *A. cruentus* groups and long-distance dispersal of the crop, potentially introducing amaranths as they migrated. More archaeological studies are necessary to elaborate the spread and influence of crops on local cultures and their exchange of plant resources.

Our finding of Mexico as center of domestication underscores the importance of this region for early agriculture and as a hotspot for crop domestication. Moreover, we cannot discard that Spanish Conquest may have influenced the grain amaranths domestication process (Sánchez-del Pino *et al*., 2025) by affecting disruption and subsequent introgression between cultivated and wild forms, which may also have contributed to the present genetic patterns. Therefore, we propose that multiple domestication scenarios should be considered to explain the observed diversity and introgression patterns in *A. cruentus*.

### Genetic structure and diversity reveal the history of *Amaranthus cruentus*

Domestication can lead to a decline in genetic diversity which might be particularly strong when populations are structured in smaller subpopulations. The subgrouping of *A. cruentus* in our analysis suggested such a geographic population structure. As mentioned previously, the region between south Mexico and Guatemala is considered as the center of domestication of *A. cruentus* (Sauer, 1967), however the exact area of the center of domestication remained challenging.

F_ST_ values indicate that *A. cruentus* group 3 has less genetic differentiation from the putative wild relative and a higher nucleotide diversity than the other two groups (Figure 4A), suggesting that it may be the closest related to *A. hybridus.* This might indicate that either *A. cruentus* group 3 is placed in an intermediate status or it is the result of higher gene flow between *A. hybridus* and *A. cruentus* group 3. This includes mainly accessions from SNICS, thus, the high nucleotide diversity found could be due to interbreeding and strategies of crop improvement.

*Amaranthus cruentus* is cultivated nowadays primarily in Guatemala and parts of central Mexico. Sauer (1950) proposed two “races” of this species:

1. **“Common race”:** Known as a grain crop in three places in Guatemala: San Martín Jilotepeque, San Juan Sacatepéquez, and Cobán. This variety is also cultivated as an ornamental plant in various countries. Despite some variation in tepal length, the floral morphology of this race is very uniform. The seeds are both pale and dark in roughly equal proportions.
2. **“Mexican race”**: Cultivated in Acatlán (Puebla), Zimatlán (Oaxaca), Ymala (Sinaloa), and Cusihuiriachic (Chihuahua) for grain production. The inflorescences are similar to the common variety, with differences mainly in the shape of the utricle cap, appearing intermediate between *A. cruentus* and *A. leucocarpus* S. Wats. (*A. hypochondriacus*) or *A. hybridus*. All grain amaranth specimens have pale seeds, except those from Cusihuiriachic.

The three genetic groups of *A. cruentus* identified in this study with k = 5 (Figure 2B and 2C) suggested that *A. cruentus* group 1 (primarily found in central Mexico but also includes some accessions from Guatemala and other northern and southern Mexican states) might represent the Mexican race described by Sauer. *Amaranthus cruentus* group 2 is mostly from Guatemala, with some accessions from Sonora and Campeche in Mexico, potentially representing the Common race. A third group (*A. cruentus* group 3) was identified, mainly includes accessions donated by the SNICS and some scattered accessions in Mexico, possibly representing a previously undescribed variety or race. These groups are evident in PCA, Admixture and Neighbor Joining analyses (Figures 2 and 3). The morphological characterization and further studies of groups found in the present study would reveal its correlation with the races previously proposed.

### Relationship of *A. cruentus* with closely related species

To further clarify the domestication process, we analyzed the genetic relationship between *A. cruentus* and its close relatives *A. hybridus* and *A. hypochondriacus. Amaranthus hybridus* is considered the wild ancestor of all three grain amaranths, with evidence suggesting multiple independent domestication processes for *A. cruentus* and *A. hypochondriacus* in Central America and *A. caudatus* in South America, whereas the wild amaranth *A. quitensis*, native to South America is thought to have played an important role in the domestication of *A. caudatus* (Sauer, 1967; Brenner *et al*., 2000; Stetter *et al*., 2020). However, *A. quitensis* and *A. hybridus* circumscription as well as *A. hybridus* polyphyletic needs to be solved to conclude any center of domestication hypothesis (Sánchez-del Pino *et al*., 2025).

Seed color, tepal posture, and bract to utricle ratios are the three characters traditionally used to identify species belonging to the hybridus-species complex; however, morphological characterization of this complex is often difficult, this is due to the existence of a few qualitative differences between the species and most of quantitative character overlap in values (Adhikary & Pratt, 2015).

In our Neighbor Joining topology, individuals of *A. hybridus* were embedded within the *A. cruentus* and *A. hypochondriacus* clades (Figure 3). This pattern reflects the high genetic diversity and cryptic substructure previously reported for *A. hybridus* and the broader hybridus-species complex (Sauer, 1967; Kietlinski *et al*., 2013; Stetter & Schmid, 2017; Wu & Blair, 2017). Although, the two crop species are strongly genetically differentiated (Figure 4a), suggesting they probably were domesticated from distinct *A. hybridus* subpopulations (Mallory *et al*., 2008).

## CONCLUSION

We identified three genetically distinct groups within *A. cruentus*, of which, two of them occur in Mexico. Our population genetic analyses support partially Saueŕs hypothesis in the sense that *A. cruentus* occurs in Mexico as the primary center of domestication, with Guatemala representing a secondary center of diversification. *Amaranthus cruentus* group 1 is located close the Tehuacán Valley, where the oldest archaeological remains of this crop have been recovered, and *A. cruentus* group 3 is located mostly in San Luis Potosí, with some individual broadly distributed in Mexico. Both Mexican groups exhibited significant introgression with *A. hybridus* and *A. cruentus* group 3 is genetically related to the wild ancestor. These findings strongly support Mexico, rather than Guatemala, as the primary center of domestication, aligning with archaeological and historical evidence.

The genomic data further reveal three well-differentiated groups of *A. cruentus*. Group 1 corresponds to the “Mexican race”, Group 2 aligns with the Guatemalan “common race”, and Group 3 may represent an uncharacterized subgroup with higher nucleotide diversity, potentially linked to interbreeding or crop improvement efforts. Finally, comparisons with closely related species underscore the central role of *A. hybridus* as the wild progenitor of grain amaranths but also reveal that different *A. hybridus* sub-populations contributed independently to the domestication of *A. cruentus* and *A. hypochondriacus*. Additional analyses and ancient DNA could help to better understand this or alternative scenarios.

## ACKNOWLEDGEMENTS

We acknowledge CONAHCYT/SECIHTI for grant under the project FORDECYT-PRONACES-15319/2019 to ISP. We thank David Brenner (USDA Curator), Eduardo Espitia (SINAREFI), and SNICS for providing plant material used in this study. We thank Andrés Xingú for growing plants at CIC’s greenhouse and collecting plant material. We acknowledge the support by the Competence Area “Food Security” of the University of Cologne for funding a six months research stay of RAST and the support by the Deutsche Forschungsgemeinschaft (DFG, German Research Foundation) under Germanýs Excellence Strategy – EXC-2048/1 – Project ID 390686111. We furthermore thank the Regional Computing Center of the University of Cologne (RRZK) for providing computing time on the DFG-funded (Funding number: INST 216/512/1FUGG) High Performance Computing (HPC) system CHEOPS as well as support.

## DATA AVAILABILITY

Raw reads will be available on ENA upon publication. Scripts used in the analysis are available on github.

