## Supplementary Figure for "Domestication of the ancient grain *Amaranthus cruentus* L"

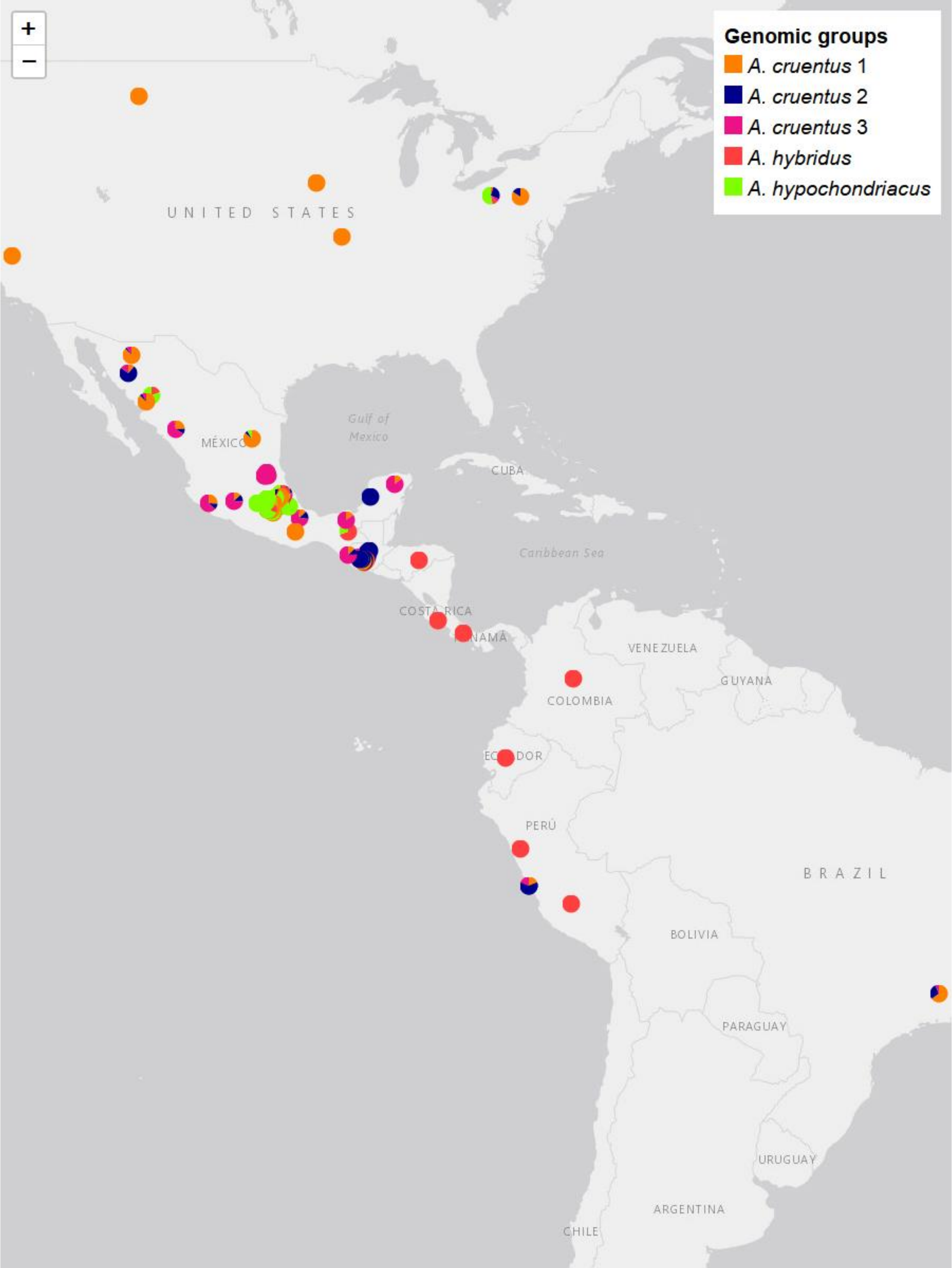


**Figure S1.** Sampling geographical distribution of individuals used in this study. Pie charts represent admixture proportions based on k = 5.

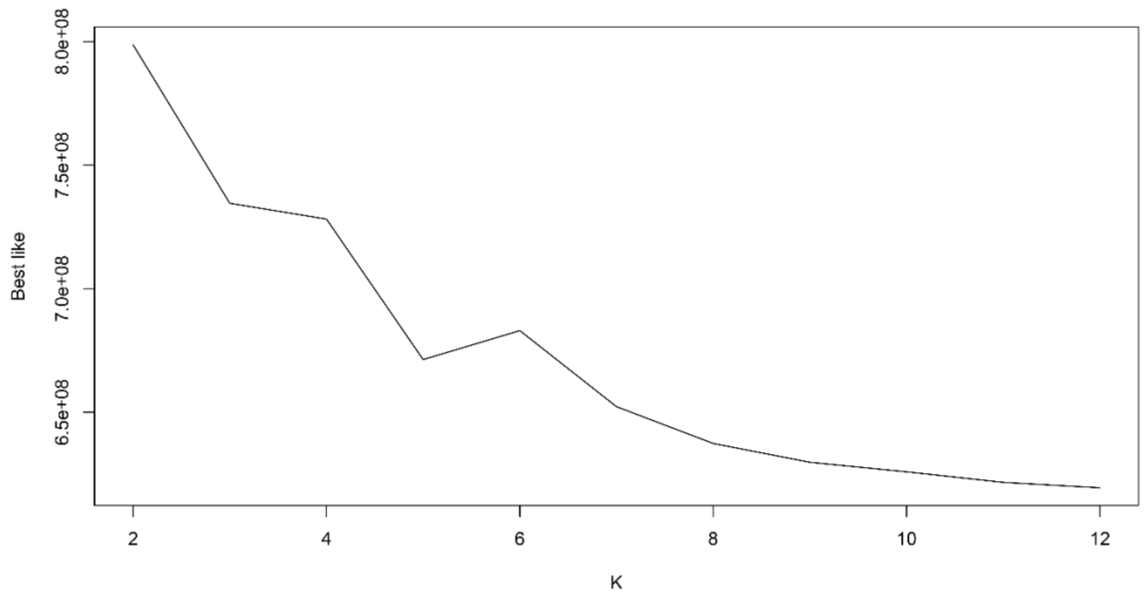


**Figure S2.** Cross validation of the admixture analysis based on elbow method. Values of k from 2 to 12 where evaluated.
